# ProteinSight: A Volumetric Deep Learning Model for Carotenoid-Binding Site Prediction

**DOI:** 10.1101/2025.10.30.685633

**Authors:** M. M. Surkov, A. Yu. Litovets, I. A. Yaroshevich

## Abstract

Carotenoproteins play essential roles across all domains of life, yet identifying them from sequence or structure remains a significant challenge due to the lack of conserved motifs. To address this gap, we present `ProteinSight`, a deep learning pipeline that identifies potential binding sites for carotenoids and related isoprenoids. Our approach, which utilizes a 3D U-Net architecture for semantic segmentation of physicochemical property maps, serves as a proof-of-concept for a new generation of structure-based protein function predictors. On a rigorously curated test set, `ProteinSight` functions as a highly sensitive and specific detector, reliably distinguishing positive from negative control proteins. Furthermore, we demonstrate its utility for hypothesis generation by predicting previously uncharacterized, plausible interaction sites on Human Serum Albumin. `ProteinSight` presents a scalable framework with the potential to aid in accelerating the discovery of novel carotenoproteins from large-scale structural data, potentially opening new avenues for functional annotation and bioengineering.

## Introduction

Carotenoids constitute one of the most extensive and functionally diverse classes of biological pigments. To date, over 1200 unique structures have been identified from a wide array of organisms, underscoring their ubiquitous distribution and fundamental importance in biological systems (1). In nature, carotenoids fulfill numerous key biological functions, acting as potent antioxidants that protect cells from oxidative stress, serving as structural stabilizers of phospholipid membranes, and participating in photosynthesis as both light-harvesting and photoprotective agents (2). Furthermore, they serve as metabolic precursors for vital molecules, including retinoids (provitamin A) (3), essential for vision, and various signaling compounds. Beyond their natural functions, carotenoids are of significant interest in medicine and biotechnology, where they are explored for their potential as therapeutic agents, including applications in oncology (4) and in overcoming multidrug resistance.

The functional repertoire of carotenoids is substantially expanded through their association with proteins to form stoichiometric complexes known as carotenoproteins. A key characteristic of these complexes is the high binding specificity conferred by a well-defined ligand-binding site. Despite the non-covalent nature of the association, the intermolecular interactions between the protein and the pigment are sufficiently precise and stable to significantly modulate the physicochemical properties of the bound carotenoid. This modulation of properties opens extensive opportunities for synthetic biology and the creation of novel biotechnological instruments. Indeed, leveraging these principles, functional molecular devices such as biocompatible photoswitches and highly sensitive thermal sensors have already been successfully engineered based on carotenoprotein scaffolds (5).

Despite the vast diversity of carotenoid chemical structures and the broad potential for directed protein modification, progress in the rational design of novel carotenoproteins is hindered by a fundamental challenge: the absence of a generalizable theory that can explain and predict the carotenoid-binding activity of proteins. The molecular mechanisms that ensure high binding specificity remain incompletely understood. To address this knowledge gap, our previous work involved a detailed statistical analysis of available atomic-resolution structures of carotenoid-protein complexes (6). This analysis revealed that carotenoid binding sites are predominantly characterized by a deep and extended hydrophobic pocket, a significant enrichment of aromatic amino acids such as tryptophan and phenylalanine, and a geometric preference for primarily perpendicular (T-shaped) similar to π-stacking interactions between the protein’s aromatic residues and the ionone rings of the carotenoid.

While such statistical analyses provide a descriptive “portrait” of a typical carotenoid-binding site, this knowledge is difficult to formalize into a deterministic predictive algorithm. The complex interplay of geometric and physicochemical features, including surface curvature, hydrophobicity, and specific intermolecular contacts, cannot be adequately captured by a simple set of rules. For practical applications, such as the de novo identification of binding sites in novel protein structures, an instrument is required that can integrate these heterogeneous features into a unified predictive model. Deep learning methodologies, particularly 3D convolutional neural networks (CNNs), have emerged as powerful tools for pattern recognition in volumetric data (7), making them ideally suited to address this challenge. These networks can learn complex, hierarchical representations of spatial and chemical patterns directly from 3D structural data without relying on predefined rules.

Here, we present `ProteinSight`, a computational pipeline based on a 3D U-Net architecture (8, 9) designed to predict the location of potential binding sites for carotenoids and related isoprenoid pigments directly from the tertiary structure of a protein. Our approach transforms a discrete atomic structure into a continuous, multi-channel volumetric representation, where each channel encodes a distinct geometric or physicochemical property of the protein’s surface and its surrounding space. By training a deep convolutional neural network on this representation, `ProteinSight` learns to perform semantic segmentation, effectively classifying voxels in 3D space based on their likelihood of belonging to a carotenoid-binding pocket. This work details the methodology for data preparation, feature engineering, and model architecture, and presents a proof-of-concept validation demonstrating the model’s ability to accurately localize known binding sites and discriminate them from other surface cavities.

## Materials and Methods

### A. Dataset Preparation

To train and validate the deep learning model, two distinct datasets were curated: a positive set comprising known carotenoid-binding and retinoid-binding proteins and a negative set containing proteins that bind other ligands but not carotenoids. All stages of this process, from initial ligand identification to the final preprocessing of structures, were fully automated through a sequence of computational scripts to ensure reproducibility.

The positive set was assembled through a two-stage procedure involving a comprehensive search for polyterpene ligands followed by the collection of their associated protein structures. First, a systematic search was performed on the official PDB Chemical Component Dictionary (CCD) (10) to identify all ligands structurally classified as polyterpenes, a class that includes carotenoids and retinoids. This was achieved by applying three specific SMARTS patterns using the `RDKit` library (11), designed to capture different types of isoprene linkages: head-to-tail, head-to-head, and tail-to-tail. Canonical SMILES were computed for each identified ligand to group synonymous PDB codes corresponding to the same molecular entity, yielding a final, unique list of ligand identifiers. Using this list, the RCSB PDB (12) API was queried to retrieve all protein structures containing at least one of these ligands. To eliminate redundancy, the resulting set of protein chains was clustered using `MMseqs2` (13) with stringent criteria of 90% minimum sequence identity and 80% minimum coverage, thereby generating a non-redundant positive dataset.

The negative set was constructed to train the model to distinguish specific carotenoid-binding sites from other concave regions on a protein’s surface, including those that accommodate different ligands. Consequently, proteins with deep, well-characterized ligand-binding pockets that do not bind carotenoids were selected as negative controls. This set was divided into two groups to ensure diversity. The general negative control group included proteins that bind common biological ligands from other chemical classes, such as nucleotides, steroids, fatty acids, and porphyrins. A random sample of up to 300 structures per ligand class was taken from the PDB. The second group, serving as a more challenging control, consisted of serum albumins, which are known for their capacity to non-specifically bind a wide range of hydrophobic molecules, making them ideal for testing model selectivity. Both negative control groups underwent the same `MMseqs2` clustering procedure as the positive set to ensure non-redundancy.

All structures from both the positive and negative datasets were subjected to a uniform and standardized preprocessing pipeline using the `pdb2pqr` software package (14). This critical step ensured the physicochemical consistency required for subsequent feature generation. The procedure involved adding hydrogen atoms, assigning protonation states to titratable residues corresponding to a physiological pH of 7.4, and assigning partial charges and van der Waals radii to each atom based on the AMBER force field (15). All crystallographic water molecules were removed. The output of this stage was a `.pqr` file for each protein structure, containing complete atomic coordinates, charges, and radii, which served as the input for the subsequent transformation into multi-channel feature fields.

To ensure a fair and rigorous evaluation of the model’s generalization capability, the dataset was partitioned into *training, validation*, and *test* sets using a strict, cluster-based splitting strategy. Operating on clusters rather than individual protein structures prevents data leakage, where a model could be tested on a protein nearly identical to one it has seen during training. First, unique cluster identifiers from both the positive and negative datasets, as generated by `MMseqs2`, were aggregated. This combined list of cluster IDs was then randomly shuffled and split into training (80%), validation (10%), and test (10%) sets. Consequently, all protein chains belonging to a single cluster were exclusively assigned to only one of these sets. This approach guarantees that the model is validated and tested on protein families entirely unseen during training, providing an unbiased assessment of its ability to generalize rather than to memorize.

### B. Voxelization and Feature Engineering

To convert the atomic protein structure into a format suitable for analysis by a convolutional neural network, a multi-step process was implemented, encompassing the definition of a computational grid, the calculation of feature fields, and their subsequent normalization.

#### B.1. Grid Definition and Setup

The space surrounding each protein structure was discretized into a regular three-dimensional grid composed of cubic voxels with dimensions of 0.5 Å × 0.5 Å × 0.5 Å. The voxel size of 0.5 Å was chosen as a balance between capturing fine atomic-level detail and maintaining computational tractability for processing large datasets. The boundaries of this grid were determined automatically for each structure by first computing the minimum and maximum coordinates of all protein atoms and then adding a padding of 12.0 Å to all sides. This padding ensures that the feature fields decay smoothly to zero at the grid boundaries, thereby preventing edge effects during subsequent convolution operations.

#### B.2. Field Calculation Method

Continuous feature fields were generated using a method wherein each atom *i* contributes to the feature value *P* at the center of each voxel *v* according to a Gaussian function. The general formulation is:

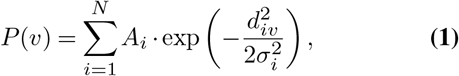

where *N* is the total number of atoms in the protein, *A*_*i*_ is the amplitude determined by the specific physicochemical property of atom *i* (e.g., its partial charge or hydrophobicity), *d*_*iv*_ is the Euclidean distance between the center of voxel *v* and the center of atom *i*, and *σ*_*i*_ is the standard deviation of the Gaussian function, which modulates the spatial influence of atom *i*. All computations were performed on a GPU using the `Numba` and `CuPy` libraries to maximize performance.

#### B.3. Description of Feature Channels

A total of eight feature channels (Figure 1, 1-8) were generated, categorized into two groups. The selection of these eight feature channels was guided by our previous statistical analysis of carotenoid-binding sites (6), which identified the importance of geometry, hydrophobicity, and aromaticity, supplemented by established descriptors of intermolecular interactions.

**Fig. 1.**
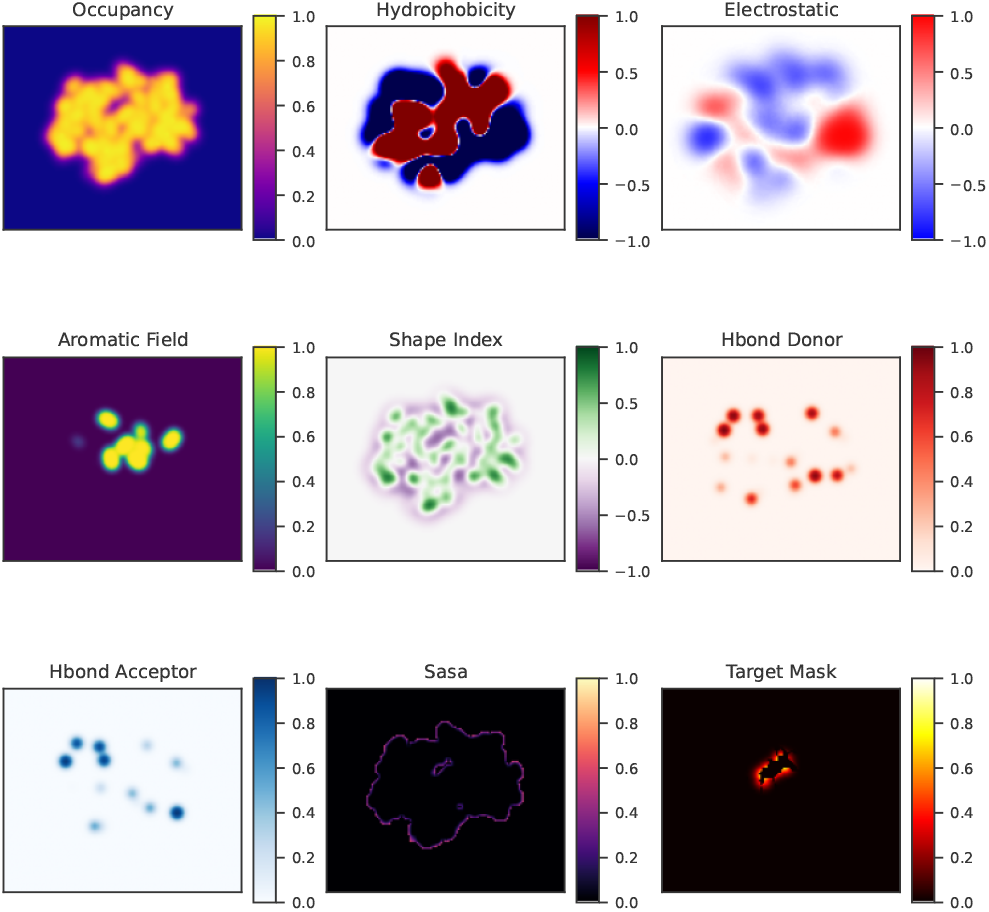
Visualization of the 8 input feature channels and the target mask for a representative red carotenoid protein (PDB ID: 5FCY). Each panel shows a 2D cross-section through the 3D tensor at the center of the binding site. These multi-modal data represent the input to the 3D U-Net model.

### A. Geometric Features

1. **Occupancy (Figure 1, 1)**. This channel describes the molecular shape. The amplitude *A*_*i*_ was set to 1 for all atoms. The standard deviation *σ*_*i*_ was made proportional to the atomic size, defined as *σ*_*i*_ = 0.9 × *R*_*i*_, where *R*_*i*_ is the van der Waals radius of atom *i* from the AMBER force field. The resulting field, *V*_occ_, was normalized using a hyperbolic tangent function to compress values into the range [−1, 1]:

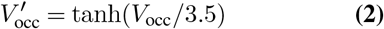
2. **Shape Index (Figure 1, 2)**. This channel characterizes the local surface curvature. It was computed as the negative of the Laplacian of the Gaussian-smoothed occupancy field (LoG). The sign was inverted so that concave pockets, which are the target regions, would have positive values. A Gaussian kernel with a standard deviation of 1.0 Å was used. The resulting field, *V*_SI_, was normalized as:

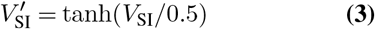
3. **Solvent Accessible Surface Area (SASA) Proxy (Figure 1, 3)**. This channel estimates the degree of solvent exposure of the protein surface. The calculation involved three steps: first, a binary mask of the protein was created; second, this mask was dilated using a spherical structuring element with a radius of 1.4 Å (approximating a water probe); third, for each voxel on the surface of the dilated mask, the number of its 26 nearest neighbors lying outside the mask was counted. This count, ranging from 0 to 26, was then divided by 26 to normalize the values to the range [0, 1].

**Fig. 2.**
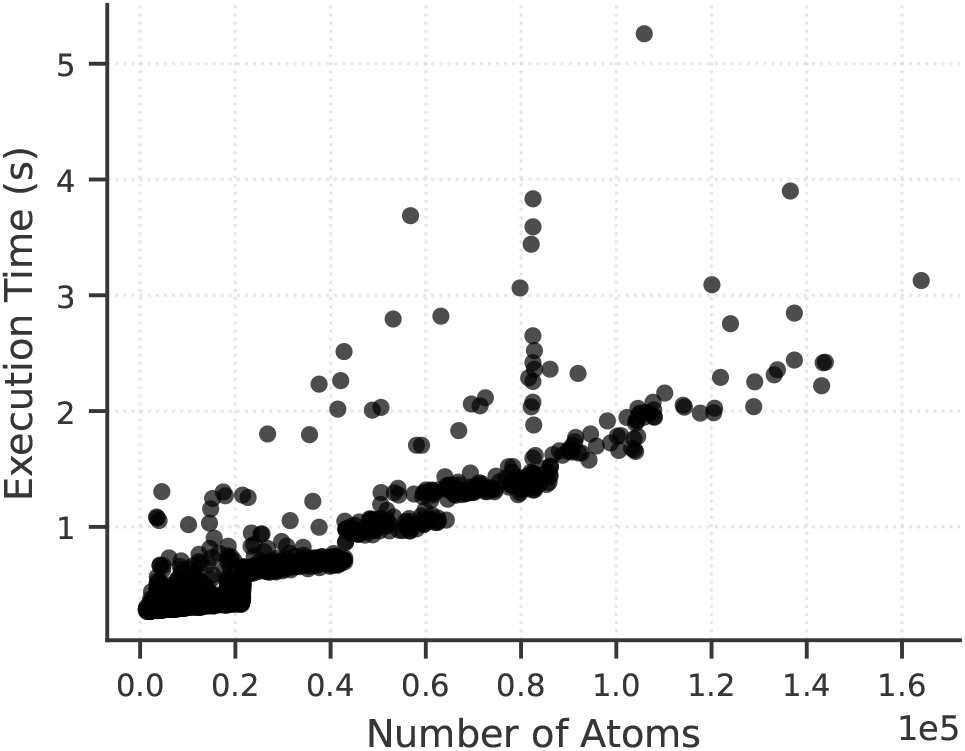
Performance of the feature generation pipeline. The execution time required to compute the full 8-channel feature field scales in a near-linear fashion with the total number of atoms in the protein. This result empirically confirms the O(N) complexity of the implemented atom-centric algorithm, demonstrating its scalability for large protein systems.

**Fig. 3.**
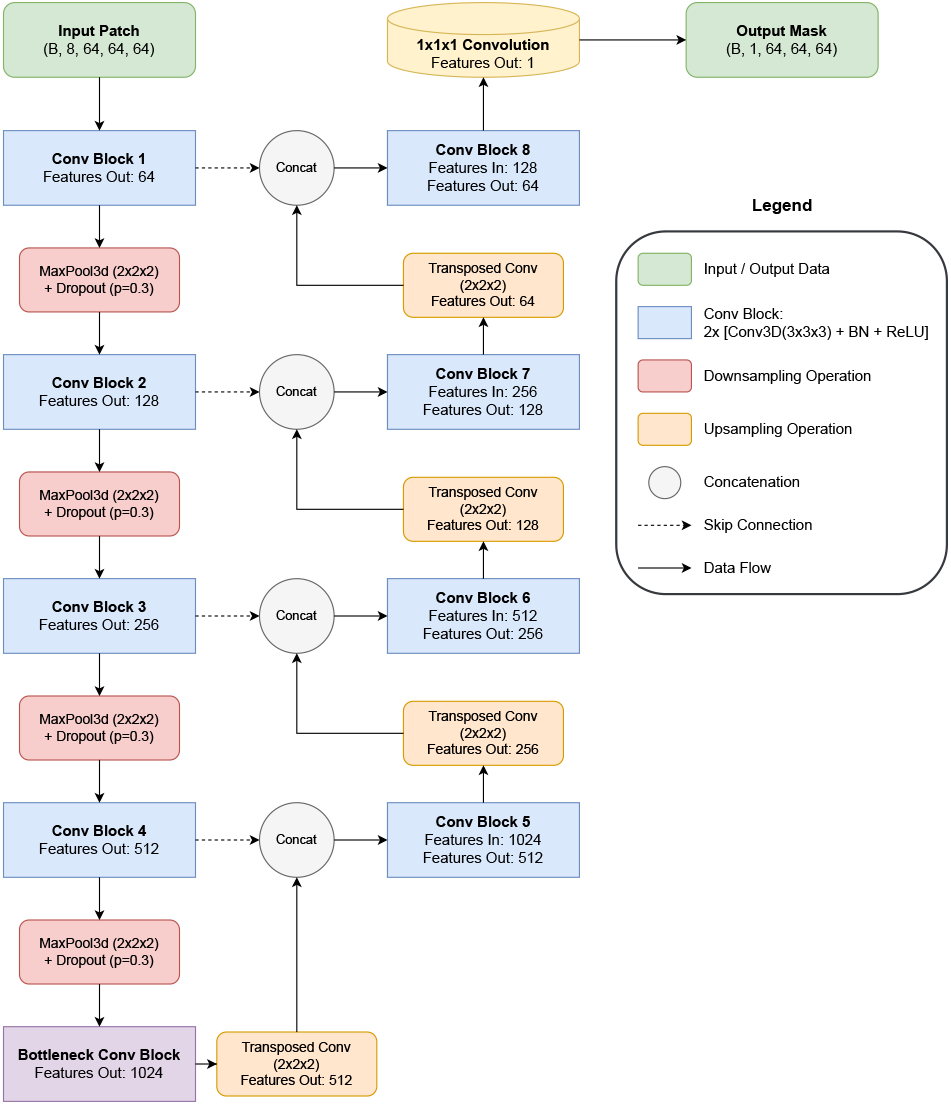
The architecture of the 3D U-Net model used for binding site segmentation. The model employs a symmetric encoder-decoder structure. The encoder path (left) consists of four blocks, each containing two sequential 3×3×3 convolutions followed by batch normalization and a ReLU activation. Downsampling is performed via 2×2×2 max-pooling with a subsequent dropout layer (p=0.3). The decoder path (right) symmetrically reconstructs the spatial resolution using 2×2×2 transposed convolutions for upsampling. Skip connections (dashed arrows) concatenate high-resolution feature maps from the encoder with the upsampled features in the decoder. A final 1×1×1 convolution reduces the feature map to a single-channel output representing the voxel-wise binding site probability.

### B. Physicochemical Features

For these channels, a constant standard deviation was used for the Gaussian function to ensure that their spatial influence was independent of atom size.

4. **Hydrophobicity (Figure 1, 4)**. The amplitude *A*_*i*_ was set to the hydrophobicity value of its parent amino acid residue according to the Kyte-Doolittle scale (16). A fixed standard deviation of *σ* = 2.0 Å was used. The field was linearly scaled from the original scale range [-4.5, 4.5] to the target range [-1, 1].
5. **Electrostatic Potential (Figure 1, 5)**. The amplitude *A*_*i*_ was set to the partial charge of atom *i* from the AMBER force field. The standard deviation was *σ* = 3.5 Å. The field, *V*_elec_, was normalized using:

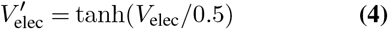
6. **H-bond Donor Potential (Figure 1, 6)**. The amplitude *A*_*i*_ was set to 1 if the atom is a hydrogen bond donor and 0 otherwise. A small standard deviation of *σ* = 1.2 Å was used to ensure a highly localized signal. The resulting field, *V*_don_, was normalized as:

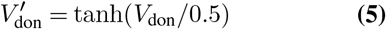
7. **H-bond Acceptor Potential (Figure 1, 7)**. This was analogous to the donor channel, with *A*_*i*_ = 1 for acceptor atoms and 0 for others, and *σ* = 1.2 Å. Normalization of the field *V*_acc_ was performed similarly:

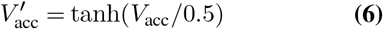
8. **Aromaticity (Figure 1, 8)**. The amplitude *A*_*i*_ was set to 1 if the atom belongs to an aromatic ring of a PHE, TYR, or TRP residue, and 0 otherwise. A standard deviation of *σ* = 1.2 Å was used to localize the signal to the rings. The field, *V*_aro_, was normalized as:

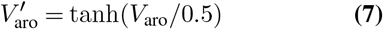

Upon completion of this process, each protein structure was represented as an 8-channel, floating-point 3D tensor, ready for input into the 3D U-Net model.

**Fig. 4.**
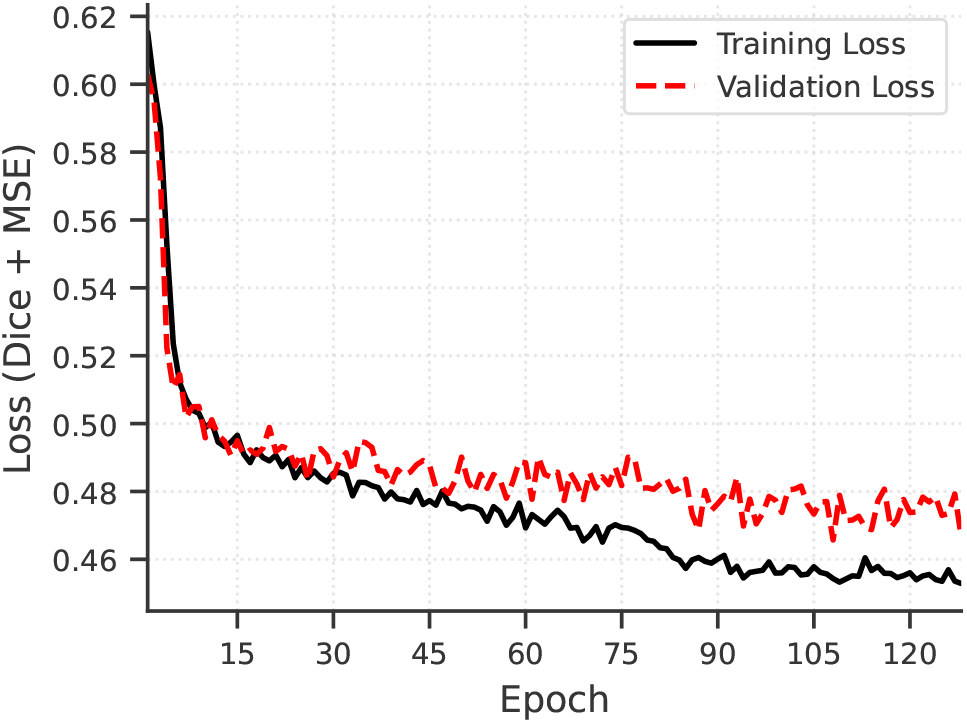
Training and validation loss curves for the model over 128 epochs. The consistent decrease and close tracking of both curves indicate successful model convergence without significant overfitting.

**Fig. 5.**
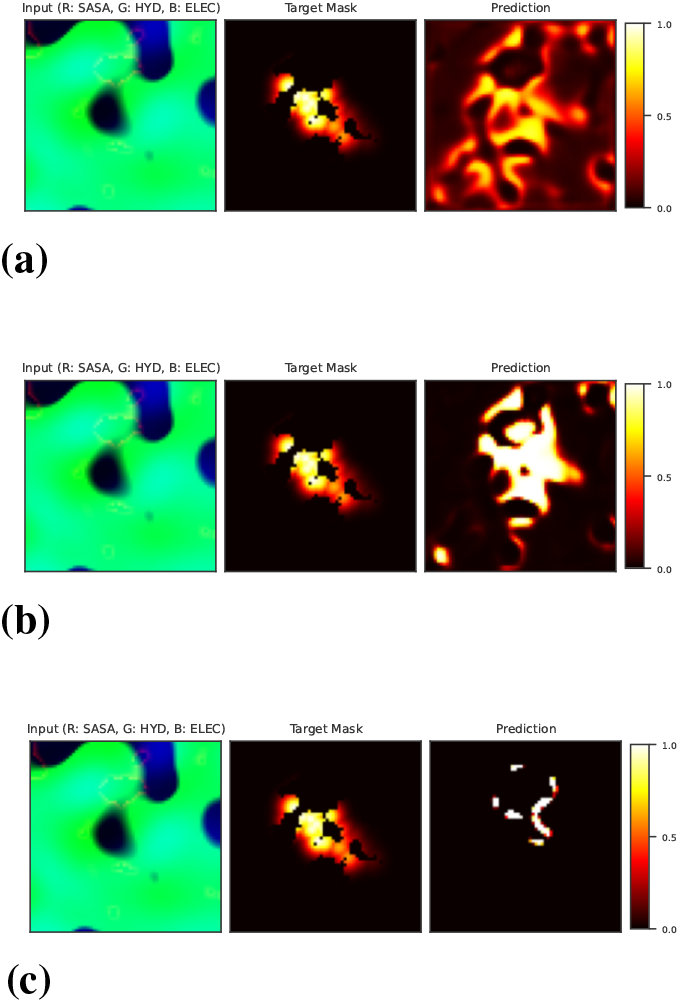
Evolution of the model’s prediction for rhodopsin (PDB ID: 1LN6) during training. (a) At an early epoch, the model predicts a diffuse region. (b) After several epochs, the prediction begins to localize. (c) In the late epoch, the prediction is sharply focused on the correct binding site. This demonstrates the model’s progressive learning and refinement.

**Fig. 6.**
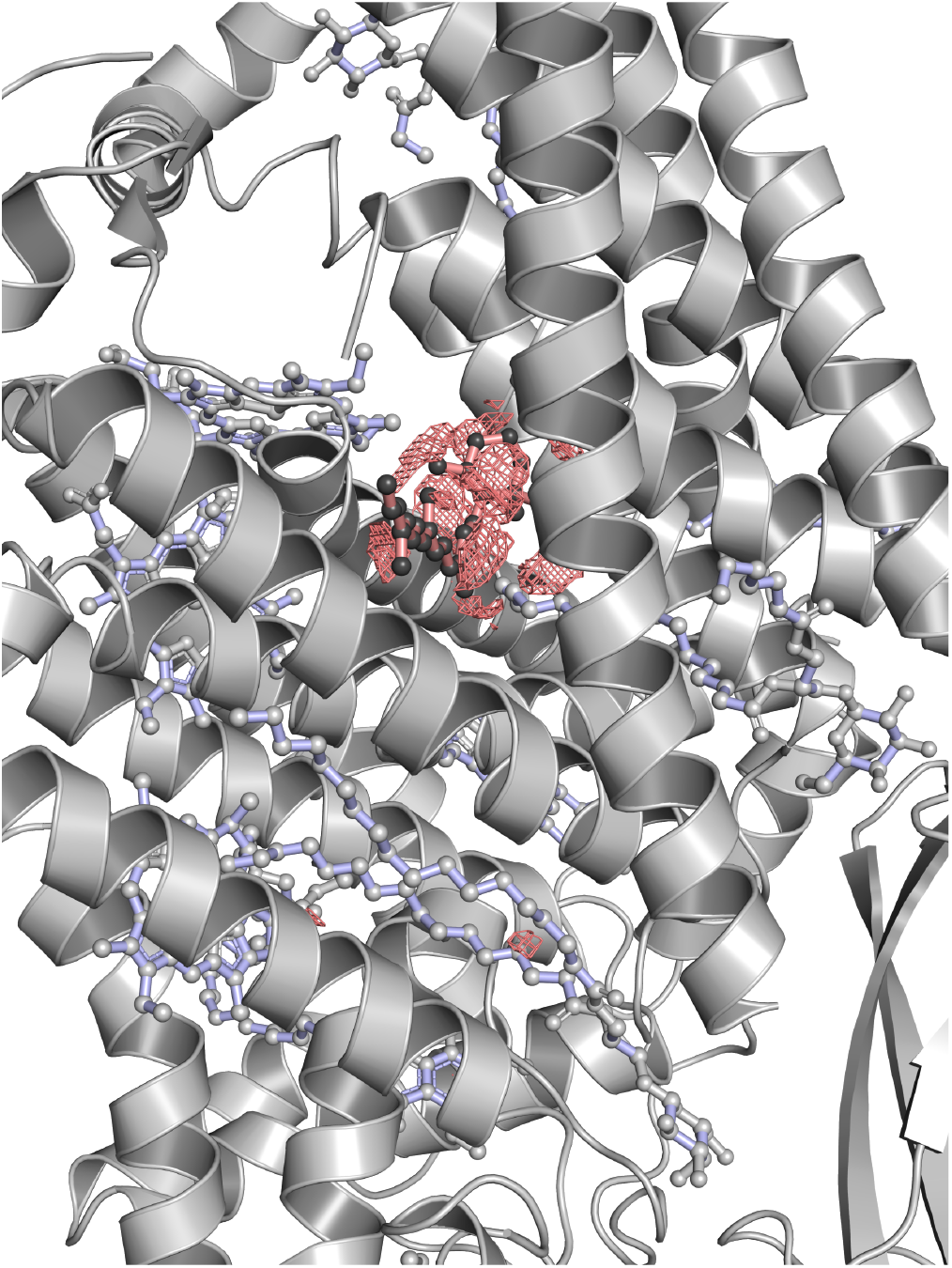
Final prediction on a positive control case, cytochrome b6f (PDB ID: 1Q90). The predicted binding site is shown as a red mesh isosurface (probability > 0.75), which accurately encapsulates the experimentally determined position of the bound beta-carotene ligand shown in ball-and-stick representation with dark gray balls and salmon sticks. The protein is depicted as a white cartoon. Other ligands are shown in ball-and-stick representation with light gray balls and light blue sticks. It is clearly noticeable that the model has accurately predicted the correct site ignoring other non-carotenoid ligands.

**Fig. 7.**
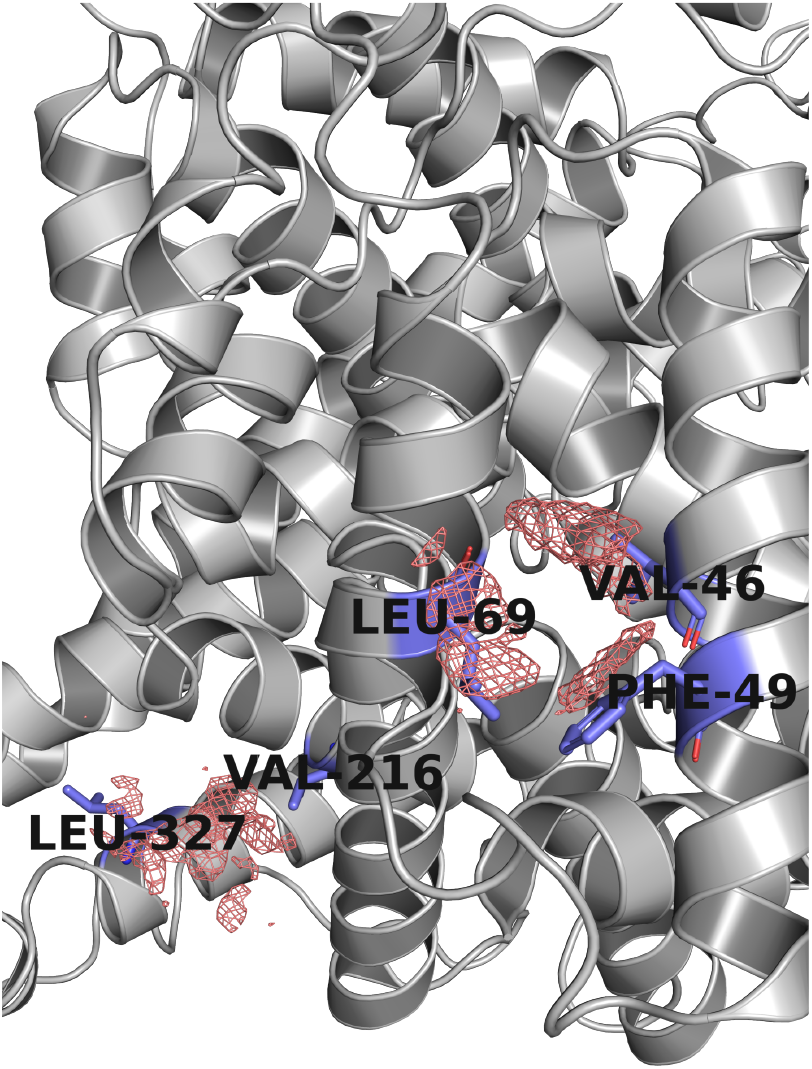
Prediction of potential carotenoid interaction sites on Human Serum Albumin (PDB ID: 1AO6). The model identifies two distinct high-probability regions (red mesh, probability > 0.5), which are not canonical binding tunnels but rather hydrophobic surface patches. These predicted sites are formed by a collection of hydrophobic and aromatic residues (shown as blue sticks), including VAL-46, PHE-49, LEU-69, VAL-216, and LEU-327, providing a structural hypothesis for the experimentally observed, non-specific binding of carotenoids to albumin. The protein is shown as a gray cartoon.

### C. High-Performance Field Generation

A key computational challenge in our pipeline is the transformation of discrete atomic coordinates into dense, three-dimensional feature fields. A naive, or “voxel-centric,” approach to this task involves iterating through each voxel in the 3D grid and summing the contributions from every protein atom. Such an algorithm has a computational complexity of *O*(*V* × *N*), where *V* is the total number of voxels and *N* is the number of atoms. For a typical globular protein, its volume, and therefore *V*, scales approximately proportionally with the number of atoms (*V* ∝ *N*), rendering the complexity of the naïve approach quadratic: *O*(*N* ^2^). This scaling makes it computationally prohibitive for processing large proteins or entire datasets.

To overcome this limitation, we implemented a significantly more efficient atom-centric algorithm. The core principle of this approach is to invert the computational logic: instead of iterating through voxels, the algorithm iterates through atoms. For each atom, its Gaussian-based contribution is computed and “splatted” onto a limited local subgrid of surrounding voxels, as its influence decays exponentially to zero at greater distances. Consequently, a roughly constant number of operations is performed for each of the *N* atoms, irrespective of the total protein size. This reduces the computational complexity to be linear with respect to the number of atoms: *O*(*N*).

To quantitatively evaluate the performance of this implementation, a benchmark was conducted on a set of proteins of varying sizes. As shown in Figure 2, the execution time required to compute the full 8-channel field (Y-axis, in seconds) scales in a near-linear fashion with the total number of atoms in the protein (X-axis). This result empirically confirms the *O*(*N*) complexity and demonstrates the high scalability of the method, enabling the efficient processing of even very large protein complexes containing hundreds of thousands of atoms in a matter of seconds. This performance was achieved by leveraging modern graphics processing units (GPUs). The implementation was developed in Python using the `Numba` and `CuPy` libraries. `CuPy` provided a NumPy-like interface for direct tensor operations on the GPU, while `Numba` was used for just-in-time (JIT) compilation of custom CUDA kernels, facilitating a highly optimized, parallelized loop over atoms for the splatting procedure.

### D. Model Architecture: 3D U-Net

For the task of semantic segmentation—that is, the voxel-wise classification of the input 3D tensor into “binding site” and “non-binding site” classes—a 3D U-Net architecture was selected and implemented (Figure 3). This architecture has demonstrated robust performance in the analysis of volumetric data, such as medical imagery, and is ideally suited to our objective of transforming an 8-channel tensor describing a protein into a single-channel probability map corresponding to the ligand’s position. The architecture is characterized by its symmetric, U-shaped structure, which comprises three primary components: a contracting path (encoder), an expansive path (decoder), and skip connections between them.

#### The Encoder Path

The encoder path is designed to extract a hierarchy of contextual features from the input patch. As data propagates deeper into the network, the encoder progressively reduces the spatial resolution (depth, height, width) while increasing the channel depth, which corresponds to the number of learned features. This process allows the model to aggregate information and recognize complex patterns at multiple scales, from local atomic arrangements to the overall shape of a binding pocket. The encoder consists of four sequential blocks. Each block includes two consecutive 3D convolutions with a 3 × 3 × 3 kernel and padding, each followed by batch normalization for training stability and a Rectified Linear Unit (ReLU) activation function to introduce non-linearity. A 2× 2× 2 max-pooling operation then downsamples the spatial dimensions of the tensor by a factor of two. A dropout layer with a probability of 0.3 is applied after pooling for regularization. At each successive block, the number of feature channels is doubled, progressing from 64 to 512.

#### The Decoder Path

The decoder path is tasked with reconstructing the original spatial resolution to produce a precise segmentation map, utilizing the high-level features extracted by the encoder. It learns to localize these features, effectively determining the precise spatial location of the patterns identified in the contracting path. The decoder is symmetric to the encoder and also consists of four blocks. Each block begins with an upsampling step performed by a 3D transposed convolution with a 2 × 2× 2 kernel and a stride of two, which doubles the spatial dimensions and halves the number of feature channels. The upsampled feature map is then concatenated with the corresponding feature map from the encoder via a skip connection. This combined tensor is then processed by two consecutive 3D convolutions, analogous to those in the encoder blocks.

#### Skip Connections

Skip connections are a crucial element of the U-Net architecture, linking the output of each encoder level to the input of the corresponding symmetric decoder level. They serve two critical functions. First, they preserve high-frequency spatial information that is lost during the pooling operations in the encoder, enabling the decoder to reconstruct segmentation masks with sharp, accurate boundaries. Second, they provide a shorter path for gradient flow during backpropagation, mitigating the vanishing gradient problem and facilitating the training of deeper networks. At each level of the decoder, the output from the transposed convolution is concatenated along the channel dimension with the output from the corresponding encoder block before being passed to the subsequent convolutions.

#### Output Layer

Finally, after the last decoder block, a final 1 × 1 × 1 convolution is applied. This layer projects the 64-channel feature tensor to a single-channel output tensor of the same spatial dimensions as the input patch. Each value in this final tensor represents a logit, or the unnormalized log-probability, that the corresponding voxel belongs to a carotenoid binding site.

### E. Training and Inference

#### E.1. Training Procedure

The model was trained using a supervised learning paradigm, wherein it learns to transform the multi-channel field representation of a protein into a probability map corresponding to the location of a binding site. To facilitate this, a target mask was generated for each protein in the positive training set (Figure 1, 9). This mask, a 3D tensor of the same dimensions as the input feature fields, was created by voxelizing the native ligand found in the crystal structure. Each heavy (non-hydrogen) atom of the ligand was represented by a Gaussian function with a fixed standard deviation (*σ*_*i*_ ≈ 2.12 Å). The value of each voxel in the target mask was assigned the maximum value from the Gaussian distributions of all nearby ligand atoms, resulting in a continuous “soft” mask. Crucially, this target was only generated within the pre-calculated protein volume mask. This method produces a volumetric, rather than surface-based, representation of the true binding site that provides a smoother learning objective for the model.×

The model was trained on 3D patches of size 64 × 64× 64 voxels extracted from the full protein feature maps. To address the severe class imbalance and provide the model with maximally informative examples, a sophisticated balanced sampling strategy was implemented. At each training iteration, the data loader generates a positive example with 50% probability and a negative example with 50% probability. For a **positive example**, a protein is randomly selected from a positive-set training cluster. A central coordinate is chosen from within the known binding site, and a random spatial jitter of up to ±32 voxels is applied along each axis. This jittering forces the model to recognize the binding site’s features regardless of its position within the patch—whether centered, at an edge, or partially occluded—thereby enhancing the model’s robustness and focus on local textural patterns over simple positional memorization.

For a **negative example**, two scenarios are chosen with equal likelihood. A **“hard negative”** example is generated by selecting a protein from a positive-set cluster but sampling a patch centered on a surface coordinate guaranteed to be distant from the true binding site. This teaches the model to perform contextual discrimination, distinguishing the target site from other cavities on the same protein. An **“easy negative”** example is generated by sampling a patch from a random surface coordinate of a protein from a negative-set training cluster. This exposes the model to generic “background” protein surfaces that lack the target site. This multi-faceted sampling scheme ensures that the model learns to identify the specific signature of a carotenoid binding site while effectively ignoring a diverse range of non-binding surface topographies.

Given the extreme class imbalance, where positive-class voxels constitute a minute fraction of the total volume, a weighted binary cross-entropy loss with logits (`BCEWithLogitsLoss`) was used as the loss function. A high weight (`pos_weight=99.0`) was assigned to the positive class, thereby penalizing false negative predictions 99 times more severely than false positive predictions. This forces the model to prioritize the correct identification of the rare but critical positive-class voxels. The Adam optimizer (17) was used with an initial learning rate of 10^*–*4^ and L2 regularization (`weight_decay=`10^*–*5^). To accelerate computations and reduce GPU memory consumption, automatic mixed-precision (AMP) training was utilized (18), enabling certain operations to be performed in 16-bit floating-point precision.

### E.2. Inference Procedure

To generate a prediction for a full protein structure, a sliding window strategy was applied. The entire 3D grid representing the protein was systematically tiled with overlapping patches of size 64 × 64 × 64 voxels, using a stride of 32 voxels. The trained model was then applied sequentially to each patch, generating an individual probability map. These maps were subsequently reassembled into a single tensor matching the dimensions of the full protein grid. In regions of overlap, predictions from multiple patches were aggregated. To produce a smooth and accurate final map, a weighted averaging scheme was employed, where predictions near the center of a patch were assigned a higher weight than those near the edges, mitigating the reduced prediction reliability often observed at the boundaries of U-Net inputs. The result of this process is a continuous 3D probability map for each protein, where the value of each voxel represents its final predicted probability of belonging to a carotenoid binding site.

## Results

### F. Model Training and Convergence

The `ProteinSight` model was trained for 128 epochs. The training dynamics were monitored by tracking the loss function on both the training and validation datasets. As depicted in Figure 4, the loss values for both sets decreased consistently and converged, indicating that the model was successfully learning the underlying patterns in the data. Importantly, the validation loss closely tracked the training loss throughout the process, showing no signs of significant divergence. This convergence behavior suggests that the model architecture and the balanced training strategy were appropriate for the task, and that substantial overfitting did not occur.

The evolution of the model’s prediction during training demonstrates a progressive refinement of its focus (Figure 5). In the initial epochs, the model identifies a broad, diffuse region of high probability that corresponds generally to the hydrophobic core of the protein. However, after only a few epochs, the prediction sharpens and becomes highly localized to the precise location of the native ligand.

### G. Qualitative Prediction on a Positive Control Case

To visually assess the predictive capability of the trained model, we performed inference on the structure of cytochrome b6f (PDB ID: 1Q90), a beta-carotene-binding protein from the test set.

The final prediction, generated by the fully trained model, shows a remarkable correspondence with the experimental structure (Figure 6). The isosurface representing a high prediction probability (threshold > 0.75) accurately delineates the elongated shape of the binding pocket and aligns precisely with the position of the bound beta-carotene molecule. This result confirms the model’s ability to correctly identify and segment a known carotenoid-binding site with high spatial accuracy.

### H. Quantitative Performance on the Test Set

To provide a rigorous and unbiased assessment of the model’s predictive capabilities, we performed a quantitative evaluation on the held-out test set, comprising 320 positive (carotenoid-binding) and 155 negative protein structures. Given the nature of the task, where identifying the presence and location of a site is more critical than segmenting its exact shape, we employed a dual-metric approach. Standard voxel-level metrics assess pixel-wise overlap, while our proposed volume-based metrics (treating a site found if it overlaps with a predicted site) evaluate the model’s ability to detect binding sites as discrete objects. The aggregated results are summarized in Table 1.

**Table 1.** Summary of model performance on the held-out test set. Voxel-level metrics assess pixel-wise accuracy, while volume-based metrics evaluate the detection of binding sites as objects. Values are reported as Mean ± Standard Deviation.

| Metric | Positive Proteins (N=320) | Negative Proteins (N=155) |
| --- | --- | --- |
| **Voxel-Level Segmentation Metrics** |  |  |
| ROC AUC | 0.919 ± 0.148 | — |
| AUPRC | 0.337 ± 0.206 | — |
| DICE F1 | 0.114 ± 0.104 | — |
| Jaccard Index (IoU) | 0.064 ± 0.067 | — |
| **Volume-Based Detection Metrics** |  |  |
| **Volume Recall** (Found Target Volume / Total Target Volume) | **0.961 ± 0.146** | — |
| **Volume Precision** (TP Pred. Volume / Total Pred. Volume) | **0.890 ± 0.273** | — |
| **F1-Score** | **0.924** | — |
| **Specificity Metric** |  |  |
| Mean Prediction Value | 0.0005 ± 0.0003 | 0.0001 ± 0.0001 |

A key indicator of the model’s discriminative power is the Receiver Operating Characteristic Area Under the Curve (ROC AUC), which remains high at **0.919**. This demonstrates that the model has successfully learned to assign significantly higher prediction scores to voxels belonging to a true binding site compared to those outside of it, confirming that the underlying learned signal is strong and reliable.

However, standard segmentation metrics that penalize shape mismatch, such as DICE F1 (0.114), are predictably low. This occurs because the model operates as a precise detector, identifying the core, high-probability region of a binding site, while the target mask represents a more diffuse, Gaussian-shaped area.

A more meaningful evaluation is provided by our proposed volume-based metrics, which treat binding sites as discrete objects. The results here are particularly strong. The **Volume Recall of 0.961** indicates that the model successfully identifies over 96% of the total volume of all true binding sites across the test set. This demonstrates sensitivity: the model rarely fails to locate a binding region. Complementing this, the **Volume Precision is at 0.890**, signifying that the model’s predictions are highly specific, with almost no predicted volume being extraneous or misplaced. In essence, what the model predicts, it predicts correctly.

Crucially, the model demonstrates excellent specificity on the negative control set. The Mean Prediction Value for these proteins was extremely low (0.0001), which is effectively background noise. This confirms that `ProteinSight` does not generate false positive predictions on proteins known not to bind carotenoids. The model has learned the specific physicochemical signature of its target and remains silent in its absence. This specificity is demonstrated directly within the cytochrome b6f example (Figure 6). The structure contains several other non-carotenoid ligands located in their own distinct pockets. Despite the presence of these welldefined binding sites, the model correctly ignored them, confining the high-probability prediction exclusively to the beta-carotene. Same results were obtained while testing the model on other negative examples, such as myoglobin. The model did not predict any high probability (>0.01) regions in negative proteins, even if they contained a deep pocket for a certain ligand.

In summary, the quantitative analysis confirms that the model functions as a highly effective and precise detector of carotenoid-binding sites. While standard segmentation metrics reflect a difference in shape between predictions and targets, the more relevant object-level metrics reveal that the model is both highly sensitive (finding >96% of true site volume) and specific (with >89% of its predictions being correct). This ability to reliably pinpoint the presence and location of binding pockets makes it a powerful tool for its primary intended application: the high-throughput discovery of novel candidate carotenoproteins.

### I. Application to a Hypothetical Case: Carotenoid Binding to Human Serum Albumin

To explore the model’s potential for generating novel hypotheses, we applied it to a particularly challenging case: Human Serum Albumin (HSA, PDB ID: 1AO6). HSA is a notoriously promiscuous carrier protein known for its ability to non-specifically bind a wide range of hydrophobic molecules. While no cocrystallized structure of an albumin-carotenoid complex exists, experimental evidence suggests that albumins can sequester carotenoids from lipid environments (19, 20). This makes HSA an ideal test case to see if `ProteinSight` can identify plausible interaction sites in the absence of a canonical, evolutionarily refined binding pocket.

Unlike the negative controls, where the model remained silent, its application to HSA yielded two distinct, highconfidence predictions (Figure 7). The predicted sites are not the elongated tunnels characteristic of dedicated carotenoproteins. Instead, the model highlights well-defined hydrophobic patches on the protein surface, composed of residues such as VAL-46, PHE-49, LEU-69, VAL-216, and LEU-327. This result is significant for two reasons. First, it demonstrates that the model’s learned features are nuanced enough to detect regions with a “carotenoid-friendly” physicochemical signature, even if they do not conform to a canonical shape. Second, it provides the first structural hypothesis for where and how carotenoids might associate with albumin, suggesting specific residues that could anchor the molecule. This case study effectively showcases the utility of `ProteinSight` not only as a tool for identifying known binding sites but also as a powerful engine for generating testable hypotheses about previously uncharacterized protein-ligand interactions.

## Discussion

In this work, we have presented `ProteinSight`, a deep learning pipeline based on a 3D U-Net architecture, designed for the identification of carotenoid-binding sites directly from protein structures. By representing proteins as multi-channel volumetric maps, our approach successfully establishes a proof-of-concept for learning and predicting the complex physicochemical environments that constitute these specialized functional sites.

Our comprehensive evaluation paints a clear picture of the model’s capabilities. The qualitative analysis on cytochrome b6f (Figure 6) demonstrated a remarkable ability to localize the known binding site while correctly ignoring pockets occupied by other non-carotenoid ligands. This specificity was further supported by the quantitative metrics (Table 1), which confirmed the model’s dual strength: a high sensitivity in detecting true sites (ROC AUC of 0.919, Volume recall of 0.961) and an excellent specificity in rejecting negative control proteins (Mean Prediction Value near zero across negative set). The performance profile is indicative of a powerful site detector rather than a precise boundary segmenter. For our primary goal of high-throughput screening and discovery of novel carotenoproteins, this is the ideal outcome, as it minimizes the risk of false positive results while roughly identifying actual site locations.

The strength of `ProteinSight` lies in its ability to move beyond the descriptive power of statistical analyses (6). While previous work could identify average properties like hydrophobicity and aromatic enrichment, our deep learning model has implicitly learned to recognize the specific, non-linear spatial combination of these features that defines a functional binding site. This capability is powerfully illustrated by the application to Human Serum Albumin (Figure 7). In this challenging case, lacking a canonical binding pocket, the model did not fail but instead generated a novel, testable hypothesis, identifying specific hydrophobic surface patches as plausible sites for non-specific carotenoid association. This demonstrates the model’s potential as a hypothesis-generation engine, moving beyond simple validation to genuine discovery.

One must, however, acknowledge the limitations of the current approach, which in turn define clear avenues for future work. A key observation is the discrepancy between the model’s sharp, localized predictions and the more diffuse, Gaussian-shaped target masks used for training. This architectural behavior explains the modest performance on voxel-level segmentation metrics like DICE, as the model excels as a precise site detector rather than a boundary segmentor. A promising direction for future refinement is to explore alternative strategies for target mask generation or to adapt the loss function to better reflect the goal of site identification over exact shape replication.

Furthermore, the current model relies on static structural data and does not account for protein flexibility, which can be crucial for ligand binding. Integrating information from molecular dynamics simulations could provide a more dynamic and realistic representation of the binding process. Finally, while `ProteinSight` demonstrates high specificity against the negative controls used in this study, its performance on a broader spectrum of proteins binding other classes of longchain hydrophobic ligands remains to be explored. Future large-scale validation will be essential to fully delineate the model’s domain of applicability.

Moreover, a comprehensive benchmark against other general-purpose binding site predictors will be conducted to formally establish its performance in a broader context. The long-term vision, however, is to deploy the trained `ProteinSight` model at a large scale on proteomes predicted by high-accuracy structure models like AlphaFold3 (21). This could facilitate high-throughput, genome-wide searches for previously uncharacterized carotenoproteins, paving the way for new discoveries in biology and new scaffolds for biotechnological engineering.

## Conclusions

In this study, we have developed and validated `ProteinSight`, a novel deep learning pipeline for the identification of carotenoid-binding sites. By transforming protein structures into multi-channel volumetric maps, our 3D U-Net model successfully learns the complex physicochemical signature of these sites. Our results suggest that `ProteinSight` operates as a sensitive and specific detector, effectively distinguishing carotenoid-binding proteins from a diverse set of non-binding control proteins. Beyond validation, the model’s capacity for generating novel, testable hypotheses—as demonstrated with Human Serum Albumin—highlights its potential as a discovery tool. In conclusion, `ProteinSight` serves as proof-of-concept for a robust computational framework poised to accelerate the discovery of novel carotenoproteins from large-scale structural data, bridging a critical gap in protein functional annotation.

## Code Availability

The complete source code for data preprocessing, model training, and evaluation is openly available on GitHub at https://github.com/MacSurmak/ProteinSight.

## Author Contributions and Use of AI Tools

M.M.S. conceived the study, developed the software, performed the analyses, and wrote the manuscript. I.A.Y. supervised the project. A.Yu.L. and I.A.Y. contributed to the manuscript revision. During the software development and manuscript preparation, large language models (including Google’s Gemini 2.5 Pro) were utilized as assistive tools for code generation, debugging, and text editing to improve clarity and correctness. All final code, analyses, and conclusions were reviewed and validated by the human authors.

